# Hemodynamic transient and functional connectivity follow structural connectivity and cell type over the brain hierarchy

**DOI:** 10.1101/2022.05.06.490832

**Authors:** Kai-Hsiang Chuang, Helena H Huang, Shabnam Khorasani Gerdekoohi, Zengmin Li, Dilsher Athwal

## Abstract

The neural circuit of the brain is organized as a hierarchy of functional units with wide-ranging connections that support information flow and functional connectivity. Studies using magnetic resonance imaging (MRI) indicate a moderate coupling between structural and functional connectivity at the system level. However, how do connections of different directions (feedforward and feedback) and regions with different excitatory and inhibitory (E/I) neurons shape the hemodynamic activity and functional connectivity over the hierarchy are unknown. Here, we used functional MRI to detect optogenetic-evoked and resting-state activities over a somatosensory pathway in the mouse brain in relation to axonal projection and E/I distribution. Using a highly sensitive ultrafast imaging, we identified extensive activation in regions up to the third order of axonal projections following optogenetic excitation of the ventral posteriomedial nucleus of the thalamus. The evoked response and functional connectivity correlated with feedforward projections more than feedback projections and weakened with the hierarchy. The hemodynamic response exhibited regional and hierarchical differences, with slower and more variable responses in high-order areas and bipolar response predominantly in the contralateral cortex. Importantly, the positive and negative parts of the hemodynamic response correlated with E/I neuronal densities, respectively. Furthermore, resting-state functional connectivity was more associated with E/I distribution whereas stimulus-evoked effective connectivity followed structural wiring. These findings indicate that the structure-function relationship is projection-, cell-type- and hierarchy-dependent. Hemodynamic transients could reflect E/I activity and the increased complexity of hierarchical processing.

**Significance Statement:** The neural circuit of the brain is organized as a hierarchy of functional units with complicated feedforward and feedback connections to selectively enhance (excitation) or suppress (inhibit) information from massive sensory inputs. How brain activity is shaped by the structural wiring and excitatory and inhibitory neurons is still unclear. We characterize how brain-wide hemodynamic responses reflect these structural constituents over the hierarchy of a somatosensory pathway. We find that functional activation and connectivity correlate with feedforward connection strengths and neuronal distributions. This association subsides with hierarchy due to slower and more variable hemodynamic responses, reflecting increased complexity of processing and neuronal compositions in high-order areas. Our findings indicate that hemodynamics follow the hierarchy of structural wiring and neuronal distribution.

## Introduction

Decoding information processing over the highly interconnected brain network requires knowledge of the hierarchy organization of its structural connectivity (SC) and structure-function relationship. Elucidating how structure and function are coupled or disrupted is not only fundamental for uncovering the neural mechanisms of behaviors, cognition and disability (1, 2), but also for early diagnosis and guiding therapeutic interventions, such as neuromodulation (3). Using magnetic resonance imaging (MRI), studies in humans have revealed that SC and functional connectivity (FC) are generally coupled at the system level (4, 5). However, the FC-SC correlation reported in the literature is highly divergent, with SC generally only explaining or predicting ~30% of the variance of functional dynamics (4, 5).

This discrepancy could be partly due to the lack of critical information such as directionality, synaptic density or excitability as MRI measures are surrogate readouts of axonal connection and neural activity. Compared with gold-standard axonal projections determined by injecting tract tracers into the mouse brain (6, 7), the SC estimated by diffusion MRI is only consistent at the system level (8). The simulated resting-state FC is also less predictable without the directionality of the SC (9). Resting-state FC measured by blood oxygenation level-dependent (BOLD) functional MRI (fMRI) is highly correlated with axonal connectivity in the cortex but not subcortical areas (10–13). Despite strong and reciprocal projections between the thalamus and cortex, the FC is absent. On the other hand, strong FC is seen in regions without a direct projection, indicating an involvement of multi-synaptic connections. Nonetheless, the lack of directionality in FC analysis makes it difficult to locate the corresponding pathways. Task- or stimulus-evoked fMRI would allow the effective connectivity to be estimated, but this relies on a clear understanding of the regional hemodynamic response function (HRF) (14). Studies in humans have shown that the cortical response is variable, which could be attributed to neural/cognitive demand, vascular density/reactivity, baseline cerebral blood flow (CBF), and neurovascular coupling (15–18). However, how the hemodynamic response propagates over the brain hierarchy and what structural factors define the responses are unclear. Excitatory and inhibitory (E/I) neurons make different contributions to neurovascular coupling, and hence could shape the regional HRF (19–22). Furthermore, microstructural factors, such as cell-type-related genes and E/I neuronal density, present a gradient over the cortical hierarchy (23–25), similar to the functional gradient found in resting-state FC (26, 27). Such structurally embedded cellular organization may underpin the regional and hierarchical variations that are not accounted for by the SC.

We hypothesized that axonal projection and E/I neuronal distribution jointly shape the propagation of the hemodynamic response and functional connectivity over the hierarchy. To test this hypothesis, we focused on the well-studied somatosensory pathway. In rodents, tactile information from the head, nose and whiskers is conveyed to the primary somatosensory cortex (SSp) via the thalamocortical projection from the ventral posteromedial nucleus (VPM) in the thalamus (7, 28). The SSp sends feedforward projections to the ipsilateral sensory, motor and association cortices, to the contralateral somatomotor cortices, and to subcortical areas, including the basal ganglia and midbrain. It also projects back to the ipsilateral thalamus via the corticothalamic projection, forming a closed loop (29). Whereas the electrophysiological properties and cell-type organization of the VPM projection to the somatosensory cortex have been studied extensively (30), the structure-function relationship in high-order areas remains elusive, partly due to the limited brain coverage of neural recordings and optical imaging. fMRI is a powerful tool for mapping whole-brain activity. Activation in the somatosensory pathway has been reliably detected in anesthetized and awake mice (31–33). Nonetheless, activation in high-order areas predicted by the SC, such as the contralateral SSp and association areas, have rarely been found because of low detection sensitivity (34, 35).

Here we investigated how directed SC and E/I organizations shape BOLD responses and FC over the hierarchy of a somatosensory pathway using optogenetic and resting-state fMRI. Compared to sensory stimulation, optogenetic excitation of the VPM allows the dissection of downstream responses while avoiding task-specific effects and bypassing processing in precedent regions, such as the brainstem (36, 37). To address the sensitivity issue, we developed an ultrafast imaging protocol that depicts whole-brain hemodynamics with 80% higher sensitivity (38). Together with axonal projection and cellular distribution mapping of the mouse brain (6, 39), we found that the hemodynamic response changed with hierarchy and local E/I activity. Although FC coupled with both the SC and E/I organization, these made differential contributions during evoked and resting states.

## Results

### Ultrafast fMRI detects brain-wide activation

To selectively activate the VPM, AAV5-hSyn-ChR2(H13R)-eYFP-WPRE was stereotaxically injected into the mouse brain to express channelrhodopsin-2 (ChR2) in all neurons. To detect activation of the VPM pathway, the BOLD signal was measured by simultaneous multi-slice echo-planar imaging that covered the entire cerebrum with 0.3s temporal resolution. Fig. 1A shows that the optic fiber targeted the dorsal VPM with low distortion despite signal loss due to a susceptibility artifact around the fiber. Histology revealed good expression of ChR2 in the VPM (Fig.1B). As stimulation frequency has a strong influence on the spatial extent of the activation (40), we evaluated the effects of the stimulating light frequency and intensity in randomized orders: 4 frequencies from 1, 5, 10 to 20 Hz were delivered in 9.9s blocks, and 5 intensities from 0.3, 0.5, 1.0, 1.25 to 2.0 mW at 5Hz were delivered as 1.5s events to minimize neural adaptation in the block design (Fig. 1C).

**Figure 1.**
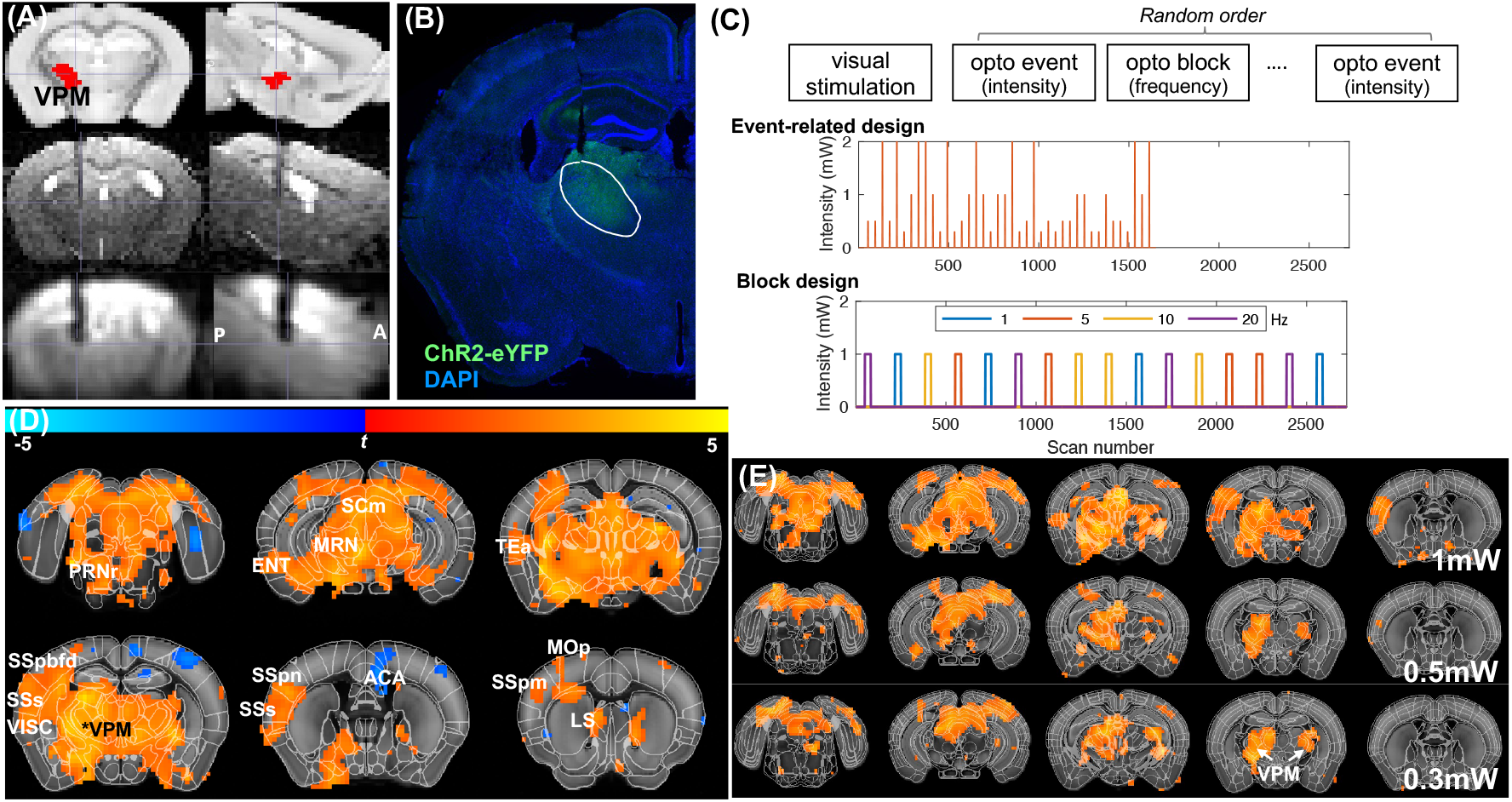
Ultrafast fMRI detects brain-wide activation under optogenetic VPM excitation. **(A)** Brain atlas (top) and structural MRI (middle) show fiber tract targeting to the VPM and the susceptibility artifact of the optic fiber in the simultaneous multislice EPI (bottom). **(B)** Histology shows ChR2 expression (green) together with DAPI (blue). **(C)** Event-related and block designs to measure the intensity- and frequency-dependent responses, respectively, under optogenetic (opto) stimulations. Each design was repeated 2-3 times in random order. **(D)** VPM excitation induces broad activation under 2mW stimulation power (p<0.05, FDR corrected). **(E)** The positive activated area reduces with lower stimulation power. See supplementary Table S2 for abbreviations of the brain regions.

After preprocessing to remove nuisance, co-registering to the Allen Brain Atlas space, and analyzing by a general linear model, our results revealed far-reaching activation in the isocortex (ISO), thalamus (TH), midbrain (MB) and hindbrain (HB), with sparse activation in the striatum (STR), pallidum (PAL), hippocampal formation (HPF) and hypothalamus of both hemispheres (Fig. 1D). With decreased stimulation intensity, the activation became restricted to the ipsilateral TH, bilateral VPM, bilateral motor-related superior colliculus (SCm) and part of the visual cortex (Fig. 1E). Similarly broad activation was seen at 1 and 5 Hz stimulations (supplementary Fig. S1A), consistent with previous studies in rats (40). Less cortical activation was seen at 10Hz. At 20Hz, the subcortical activation was reduced and more restricted to the ipsilateral TH and the SCm. Interestingly, negative BOLD responses were consistently found in the contralateral cortex at all the stimulation frequencies. This demonstrated the detection of downstream activation throughout the brain with enhanced fMRI sensitivity. Although a block design is typically used in rodent imaging for its high statistical power, strong adaptation was found even at the VPM (Supplementary Fig. S1B). Hence the following analysis focuses on the event-related data.

### Activation matches SC hierarchy

We constructed the system-level hierarchy across 350 brain regions based on axonal projections derived from the monosynaptic anterograde tract tracing data in the Allen Brain Connectivity atlas (6). SC strength was calculated as the summed axonal projection volumes, normalized by the tracer injection volume. From the VPM, we identified up to 74 target areas, distributed in the ISO, TH, MB and HB, which receive direct (first-order) projections (supplementary Fig. S2A). The strongest projections were the secondary somatosensory cortex (SSs) and the upper lip (SSpul), mouth (SSpm), nose (SSpn) and barrel field (SSpbfd) in the SSp, reticular nucleus of the thalamus (RT) and SCm of the ipsilateral hemisphere. There were also contralateral projections to the above somatosensory areas, but the projection strengths were at least 68-fold weaker. From the somatosensory cortex, much broader second-order connections could be found. For example, the SSpbfd projected to 233 regions (supplementary Fig. S2B), with strong feedforward projections to the ipsilateral caudate putamen, primary motor cortex (MOp), secondary motor cortex (MOs) and SSs, and to the contralateral barrel field (SSpbfdc), as well as feedback projections to the VPM and posterior complex of the thalamus.

The hierarchy could be better visualized by removing weak connections. As the SC follows a lognormal distribution (6), a threshold was defined based on the logarithm of the SC (supplementary Fig. S2C). By choosing the top 10% strongest connections, the hierarchy showed that the major first-order areas are in the ipsilateral somatosensory cortex (Fig. 2A). The second-order afferents included feedforward projections to other sensory and motor areas in both hemispheres and feedback projections to the ipsilateral somatosensory areas and thalamus. The third-order afferents extended to limbic and association areas, such as the anterior cingulate area. The peak amplitude of the evoked BOLD response in each region was tested and showed significant change (p<0.01 uncorrected) in >96% of the regions predicted by the SC. Compared to this structural hierarchy, up to third-order areas could be identified by fMRI.

**Figure 2.**
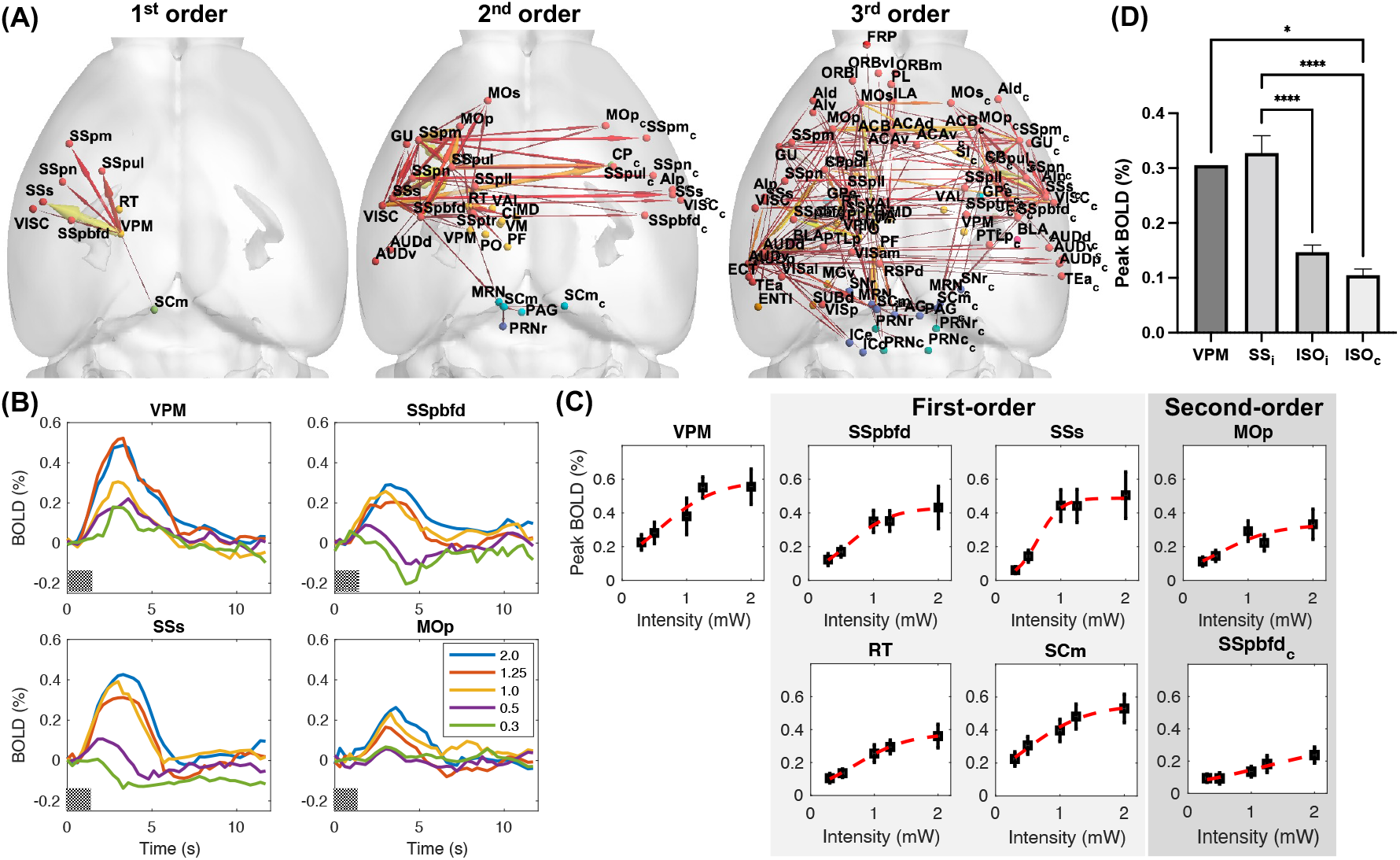
Regional BOLD responses reduce with hierarchy. **(A)** The structural hierarchy defined based on axonal projection. The 1^st^-order areas (left) receiving direct projections from the VPM, 2^nd^-order areas (middle) receiving direct projections from the 1^st^-order areas, and the 3^rd^-order areas (right) receiving direct projections from the 2^nd^-order areas, based on thresholding of the top 10% SC. **(B)** Averaged BOLD signal time-courses in the VPM, two first-order afferent areas (SSpbfd and SSs) and a second-order area (MOp) under 5 stimulation intensities (in mW). The gray bar presents the stimulation period. **(C)** The stimulation intensity-dependent responsiveness follows a sigmoid function. Error bars present the SEM. **(D)** BOLD activation amplitudes are highest in the first-order areas and decrease in the second-order isocortex (ISO) of the ipsilateral (ISO_i_) and contralateral (ISO_c_) hemispheres. *: p<0.05, ****: p<0.0001.

### Cortical BOLD responses change with hierarchy

To understand how hemodynamic responsiveness changes with hierarchy, we examined the evoked BOLD responses of different stimulation intensities (Fig. 2B). The VPM activation increased with stimulation intensity and reached a plateau at around 1.25 to 2 mW. The responses in the first-order cortical areas, such as the SSpbfd and SSs, followed a similar trend but plateaued slightly earlier. In the second-order areas, such as the MOp, the activation was weaker and had a different trend. This was more apparent when fitting the peak BOLD responses by a sigmoidal function (Fig. 2C). Compared to the sigmoid trend in the VPM, the trends in the TH and MB were highly similar (r=0.99±0.003, mean±standard error) regardless of the hierarchy. In contrast, those in the ipsilateral ISO were variable (r=0.89±0.058) and more divergent in the contralateral ISO (r=0.59±0.17). The cortical intensity response curves became more variable with hierarchy (supplementary Fig. S3), suggesting increased complexity of regional processing in higher-order areas. As the cortical activation tends to plateau above 1mW, we assessed the peak BOLD amplitude at 1mW stimulation. The BOLD amplitudes reduced with hierarchy (F_3,63_=17.21, p<0.0001), with activations in the second-order ISO being significantly weaker (p<0.0001) than that of the first-order somatosensory area (Fig. 2D). The more variable and weaker activation in high-order cortical areas may explain their poor detectability in previous studies.

Previous studies in humans showed that hemodynamic responses of different cortical areas are variable (15–17), whereas studies in rodents found similar responses (33, 41). Whether hemodynamic variation follows structural organization is unclear. Intriguingly, we found distinct BOLD responses in subcortical regions (Fig. 3A) and the cortical hierarchy. BOLD responses were similarly sharp in the TH and MB, slightly broad in ISO, but slow in the STR and PAL. In particular, biphasic responses were consistently found in the contralateral cortex but not subcortical regions. These responses were not dependent on the hierarchy as the second-order ipsilateral areas, such as the Mop, still showed a positive response (Fig. 3B). To characterize regional impulse responses, we deconvolved and fitted the BOLD signal with double gamma variate functions. Based on the hierarchy of major projections, the hemodynamic impulse responses were more variable at higher order (Fig. 3C). The time-to-rise, defined as the time when a signal increases to 30% of the peak, in the higher order areas were significantly longer than those in the first-order areas (F_2,21_=4.93, p=0.017; Fig. 3D). The time-to-positive-peak in the ipsilateral cortex also increased over the hierarchy (F_2,20_=5.45, p=0.013) from 2.48±0.14 s (mean±standard error) to 3.51±0.32 s (Fig. 3E). The responses in the contralateral cortex had shorter time-to-positive-peak but longer time-to-negative-peak than the peak timing in the ipsilateral cortex of the same hierarchy. However, these contralateral peak timings were not hierarchy dependent. These results suggest that the initial timing of the hemodynamic response could reveal hierarchical relationship.

**Figure 3.**
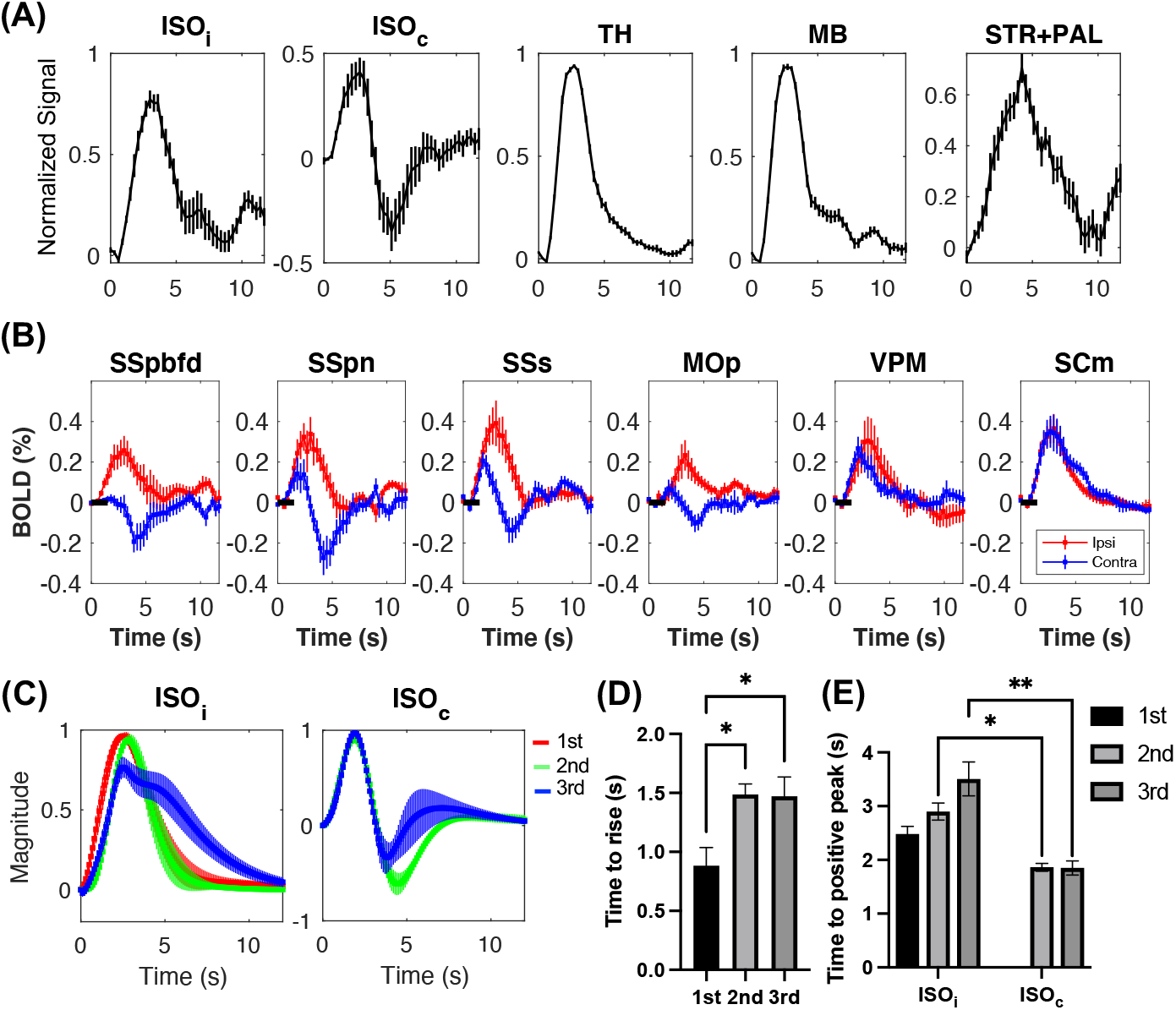
Hemodynamic response is structural organization and hierarchy dependent. **(A)** Mean evoked BOLD of the ipsilateral and contralateral ISO, TH, MB, and STR+PAL measured at 1mW stimulation. **(B)** Examples of ipsilateral and contralateral BOLD responses in the somatosensory and primary motor cortices, and in the VPM and superior colliculus. The black bar represents the stimulation period. **(C)** Fitted response functions over the three orders of major SC hierarchy in the ipsilateral and contralateral ISO. **(D)** The hemodynamic time-to-rise over the hierarchy in the ipsilateral ISO. **(E)** The time-to-positive peak over the hierarchy in the ipsilateral and contralateral ISO. *: p<0.05. Error bars represent the SEM.

Time-lag regression analysis, such as Granger causality, has been used to estimate causality. To test whether hemodynamic timing could be estimated by regression, we used crosscorrelation analysis to estimate the BOLD signal lag time with respect to the VPM. Surprisingly, the lag time did not correlate with the time-to-rise or time-to-peak, but did correlate with the peak BOLD amplitude (r=-0.4, p<0.0001; supplementary Fig. S4). The lag time did show a trend (Kruskal-Wallis statistics=6.88, p<0.05) to increase over the hierarchy in the ipsilateral cortex, likely due to hierarchy-dependent reduction of the BOLD amplitude. This indicates that although time-lag analysis may show hierarchy-dependent change, it does not reflect the actual hemodynamic timing.

### Hierarchical BOLD activation corresponds to the local field potential

Whether the hemodynamic changes over the hierarchy are due to differences in neural activity or the regional CBF and neurovascular coupling is unclear. To investigate this, we conducted intracortical field recording in three ipsilateral areas, SSs, MOp and dorsal ACA (ACAd), which correspond to primary, secondary and tertiary projection sites, respectively. With the same optogenetic stimulation at the VPM, strongly evoked local field potentials (LFPs) were observed in the SSs and MOp whereas weak responses were seen in the ACAd (Fig. 4A). The integral of the LFP magnitude showed a strong increasing trend with stimulation intensity at the SSs but was weaker at the MOp and ACAd (Fig. 4B). Comparison of the LFP with the peak BOLD signals over the range of stimulation intensities revealed a linear relationship at the SSs (R^2^=0.92, p=0.010), MOp (R^2^=0.78, p=0.047) and ACAd (R^2^=0.93, p=0.0089) (Fig. 4C). In particular, the similar slope between the LFP and the BOLD signals suggested that the neurovascular coupling of these areas are comparable. To determine the response latency, the time-to-rise of the first LFP peak was used to represent the initial neuronal population activity (Fig. 4D). The time-to-rise increased from 3.65±0.39 ms at the primary target (SSs) to 6.33±0.26 ms at the secondary target (MOp) and 9.13±0.43 ms at the tertiary target (ACAd) (F_2,11_=53.97, p=2×10^-6^). The full-width-at-half-maximum (FWHM) of the highest LFP peak was used to characterize the dispersion of population activity. The FWHM increased from 6.98±0.99 (SSs) to 8.75±0.41 (MOp) and 12.33±0.50 ms (ACAd) over the hierarchy (F_2,11_=16.82, p=0.00045).

**Figure 4.**
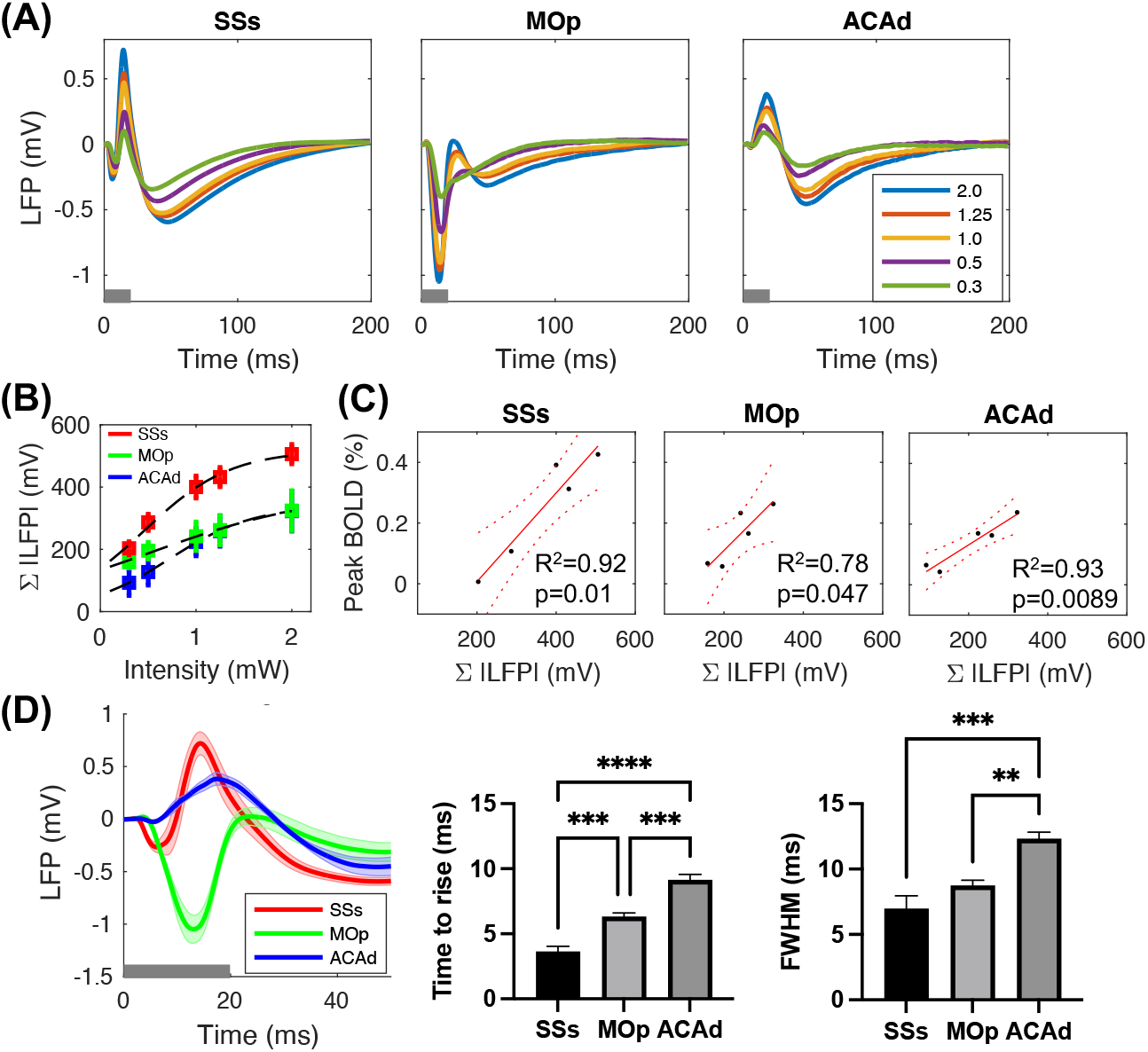
LFP over the hierarchy. **(A)** Averaged evoked potentials at the SSs (N=4), MOp (N=6), ACAd (N=4) under 5 intensities (mW) of optogenetic stimulation events (20ms pulse, 5Hz and 1.5s duration) at the VPM. **(B)** The integral of evoked potential magnitudes (Σ|LFP|) over the stimulation intensities. The dashed line is the fitted sigmoid function. **(C)** Linear regression between the LFP and BOLD signals. The dashed lines represent the 95% confidence interval. **(D)** The comparison of evoked potentials under 2mW stimulation shows the initial timing and response width differences between regions. An early negative peak followed by a high positive peak was seen at the SSs. The time-to-rise of the first peak (one-way ANOVA, F_2,11_=53.97, p=2×10^-6^) and FWHM of the highest peak (F_2,11_=16.82, p=0.00045) increased over the hierarchy. The error bars represent the SEM. **: p<0.01, ***: p<0.001, ****: p<0.0001.

Previous studies showed that lower baseline CBF can lead to larger BOLD activation amplitude (18, 42, 43). To assess the contribution of regional vascular function, the CBF was quantified using arterial spin labeling MRI. The cortical hemodynamic amplitude, time to peak and response width (FWHM) did not correlate with baseline CBF (supplementary Fig. S5A). Despite an increased variation in CBF in the high-order hierarchy (supplementary Fig. S5B), this could not explain the BOLD variations (supplementary Fig. S5C). Together, these results indicate that cortical BOLD variations reflect neural activity, rather than the regional neurovascular coupling, over the hierarchy.

### BOLD activation and FC correlate with feedforward projection

To understand how SC influences BOLD activity, we compared their relationship over thalamocortical (feedforward), corticocortical (feedforward and feedback) and corticothalamic (feedback) projections (Fig. 5A; supplementary Table S1). In the thalamocortical projection, the peak BOLD amplitudes in the targeted cortical areas highly correlated with the SC from the VPM (R^2^=0.77, p=6.4×10^-5^; Fig. 5B). However, the BOLD activation in the subcortical areas (TH, MB and HB) was not associated with feedforward from the VPM. The corticocortical feedforward from most of the somatosensory areas to the ipsilateral cortex also correlated with the BOLD activation in these second-order areas, with R^2^ ranging from 0.16 to 0.79 (Fig. 5C). However, we observed no association in the corticocortical feedforward to the contralateral cortex, except in the case of the interhemispheric somatosensory areas (R^2^=0.88, p<0.05; Fig. 5D). There was also no association in corticocortical feedback among the somatosensory areas. In the corticothalamic feedback, we found that the thalamic BOLD activation correlated with the SC from the SSpbfd and SSpm (Fig. 5E), but not the other somatosensory areas. Notably, the association with feedforward SC was not observable in the block-design data (supplementary Fig. S6). The proportionally higher BOLD response over stronger SC supports the typical assumption of neural modeling that regional activation depends on the projection strength. However, this coupling varied among projections, diminished in feedback projections, and was abolished by nonlinear effects under a longer stimulation.

**Figure 5.**
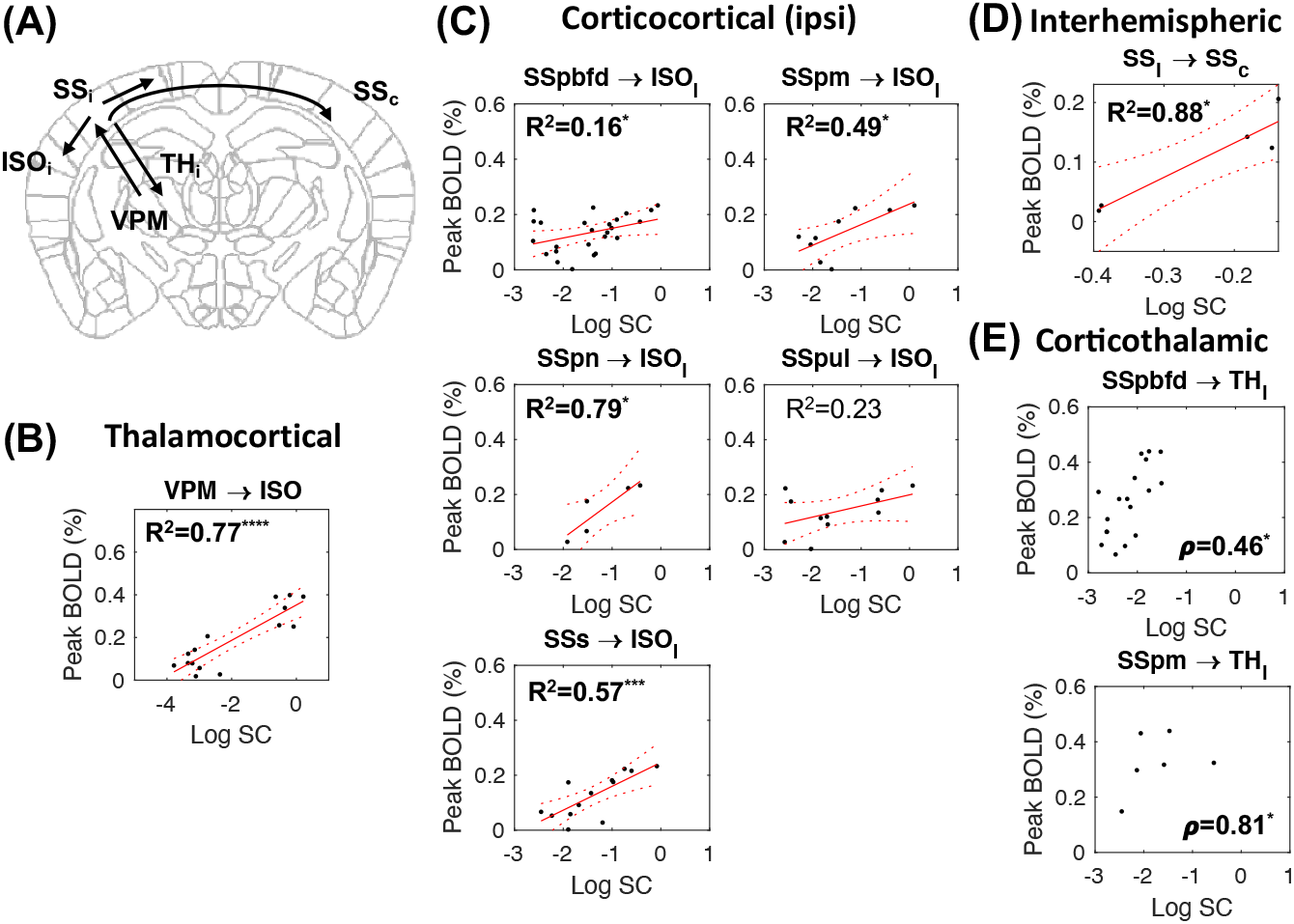
BOLD activation correlates with SC. **(A)** Diagram of thalamocortical (VPM→ISO), corticocortical (SS_i_→ISO_i_, SS_i_→SS_c_) and corticothalamic (SS_i_→TH_i_) projections. Linear regression between the Log SC and the peak BOLD amplitude at the target areas measured at 1mW stimulation in the **(B)** thalamocortical feedforward, **(C)** corticocortical feedforward from the ipsilateral somatosensory cortex (SS_i_) to the ipsilateral ISO, and **(D)** corticocortical feedforward from the ipsilateral to contralateral somatosensory cortex (SSc). **(E)** Spearman correlation between the Log SC and peak BOLD amplitude in the corticothalamic feedback from the somatosensory cortex to the ipsilateral thalamus. *: p<0.05, ***: p<0.001, ****: p<0.0001. Solid lines represent the linear fit and dashed lines represent the 95% confidence interval. See supplementary Table S1 for complete results.

To determine the FC-SC coupling, we calculated the FC by zero-lag correlation coefficients between regional time-courses under optogenetic stimulation (FC_Task_) or resting state (FC_Rest_). To inspect the relationship between regional activation, the stimulation response was not regressed from the FC_Task_. Unlike FC_Rest_, which showed strong cortical but weak subcortical connectivity, the FC_Task_ was strong within and between the TH and MB (Fig. 6A). On the other hand, the interhemispheric cortical FC_Task_ was weak due to the hemodynamic difference between hemispheres. Despite these differences, the FC_Rest_ highly correlated with FC_Task_ over the whole brain (R^2^=0.62), ISO (R^2^=0.57, p=7.9×10^-67^) and TH (R^2^=0.81, p=8.1×10^-156^).

**Figure 6.**
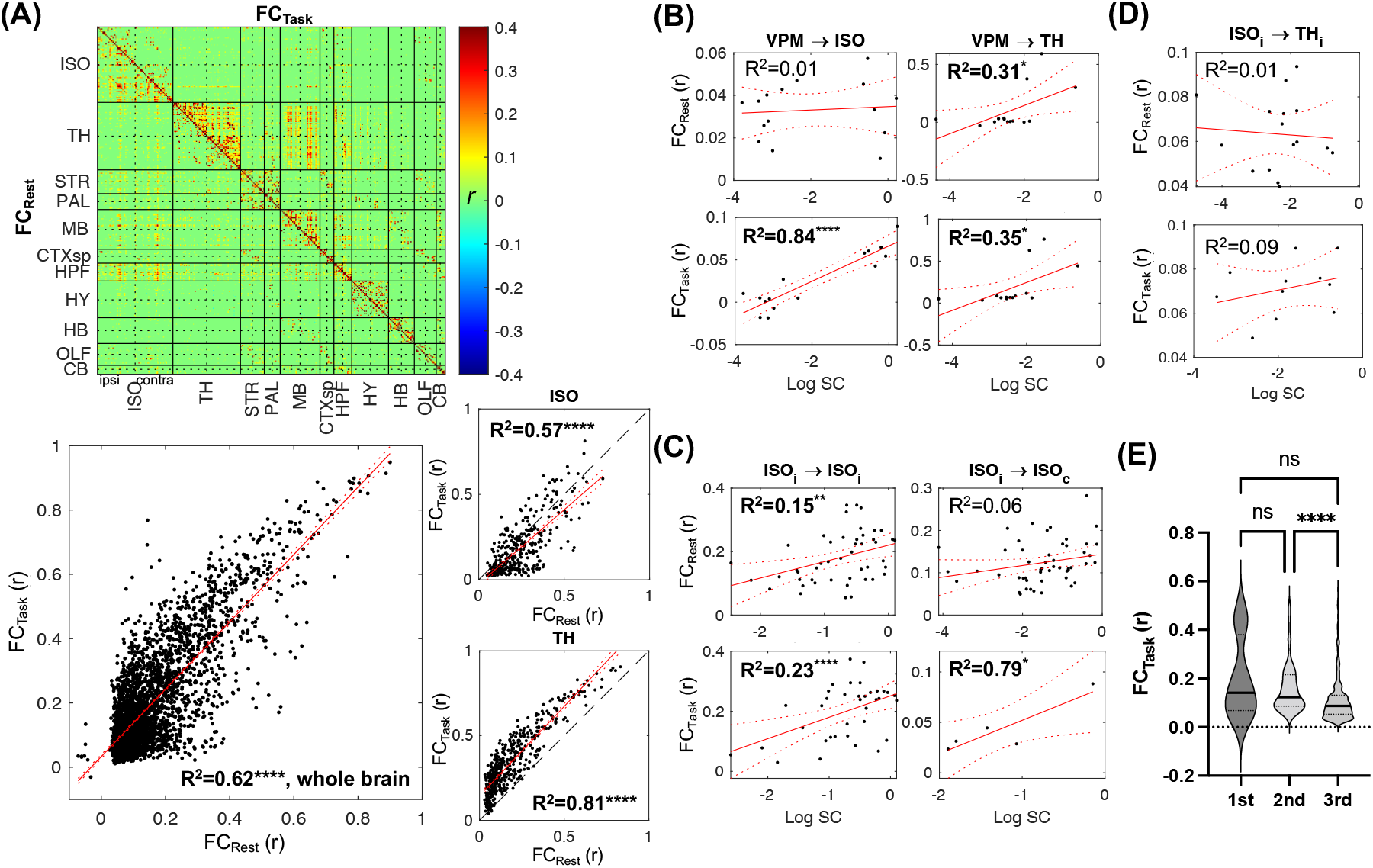
Comparison between FC_Rest_, FC_Task_ and SC. **(A)** The connectivity matrices of FC_Rest_ (lower triangle) and FC_Task_ (upper triangle) thresholded at p<0.01, and their correlation in the whole brain, ISO and TH. Linear regressions between the FCRest and Log SC (upper row) and between the FC_Task_ and Log SC (lower row) in the (**B)** thalamocortical and thalamic-thalamic feedforward, (C) corticocortical feedforward, and **(D)** corticothalamic feedback. **(E)** FC_Task_ over the hierarchy of major SC in the ipsilateral ISO. *: p<0.05, **: p<0.01, ***: p<0.001, ****: p<0.0001. Dashed lines represent the 95% confidence interval. CTXsp: cortical subplate, HPF: hippocampal formation, HY: hypothalamus, OLF: olfactory bulb, CB: cerebellum.

The FC_Task_ correlated with the SC in the thalamic (R^2^=0.35, p<0.05) and thalamocortical (R^2^=0.84, p<0.0001) feedforward from the VPM (Fig. 6B). In the cortex, the FC_Task_ correlated with the corticocortical feedforward within and between hemispheres, with R^2^ = 0.23 to 0.84 (Fig. 6C). In comparison, FC_Rest_ only correlated with SC in a few connections. However, neither FC_Task_ nor FC_Rest_ correlated with the SC in the corticothalamic feedback (Fig. 6D). There was also a slight trend towards decreased FC over the hierarchy (Fig. 6E) which could be due to weaker and more variable BOLD responses at higher order. These results show that FC_Task_, but not FC_Rest_, has moderate to high correlation with directed SC, particularly in feedforward projections.

### Cortical hemodynamics and FC associate with E/I organization

Previous studies of the spatial association between FC and gene expression pattern suggested that cellular organization and micro-circuitry may underlie this correlation (44). However, whether regional E/I neuronal composition contributes to the differences in hemodynamic response and FC is unclear. We extracted the neuronal densities of excitatory neurons and 3 major types of inhibitory interneurons that express somatostatin (SST^+^), parvalbumin (PV^+^), and vasoactive intestinal peptide (VIP^+^), respectively (39, 45). Linear regression was used to predict hemodynamic parameters, including timing, amplitude and width, based on the neuronal densities. We found that excitatory, but not inhibitory, neuronal density contributed to the time-to-peak (R^2^=0.18, p=0.0077) and FWHM (R^2^=0.11, p=0.039) of the positive BOLD responses in the cortex (Fig. 7A and supplementary Fig. S7). The time-to-rise and FWHM of the negative BOLD responses were influenced by the densities of inhibitory SST^+^ (R^2^=0.36, p=0.0037) and VIP^+^ (R^2^=0.59, p=0.00005) neurons, respectively (Fig. 7B). In contrast, the hemodynamic parameters of subcortical areas, such as the TH, did not correlate with any of the neuronal types. Interestingly, the E/I ratio, calculated by dividing the excitatory neuron density by the sum of the 3 types of inhibitory neurons, increased over the hierarchy (Kruskal-Wallis statistic=9.495, p=0.0087) due to decreased SST^+^ (F_2,48_=4.14, p=0.022) and PV^+^ (F_2,48_=4.09, p=0.023) neuronal densities over the hierarchy of major connections (supplementary Fig. S8). This partly explained the hierarchy-dependent signal variation as the cortical BOLD similarity decreased with E/I ratio (ρ=-0.28, p=0.013; supplementary Fig. S8C). These results suggest that hemodynamic characteristics may reflect underlying E/I activity.

**Figure 7.**
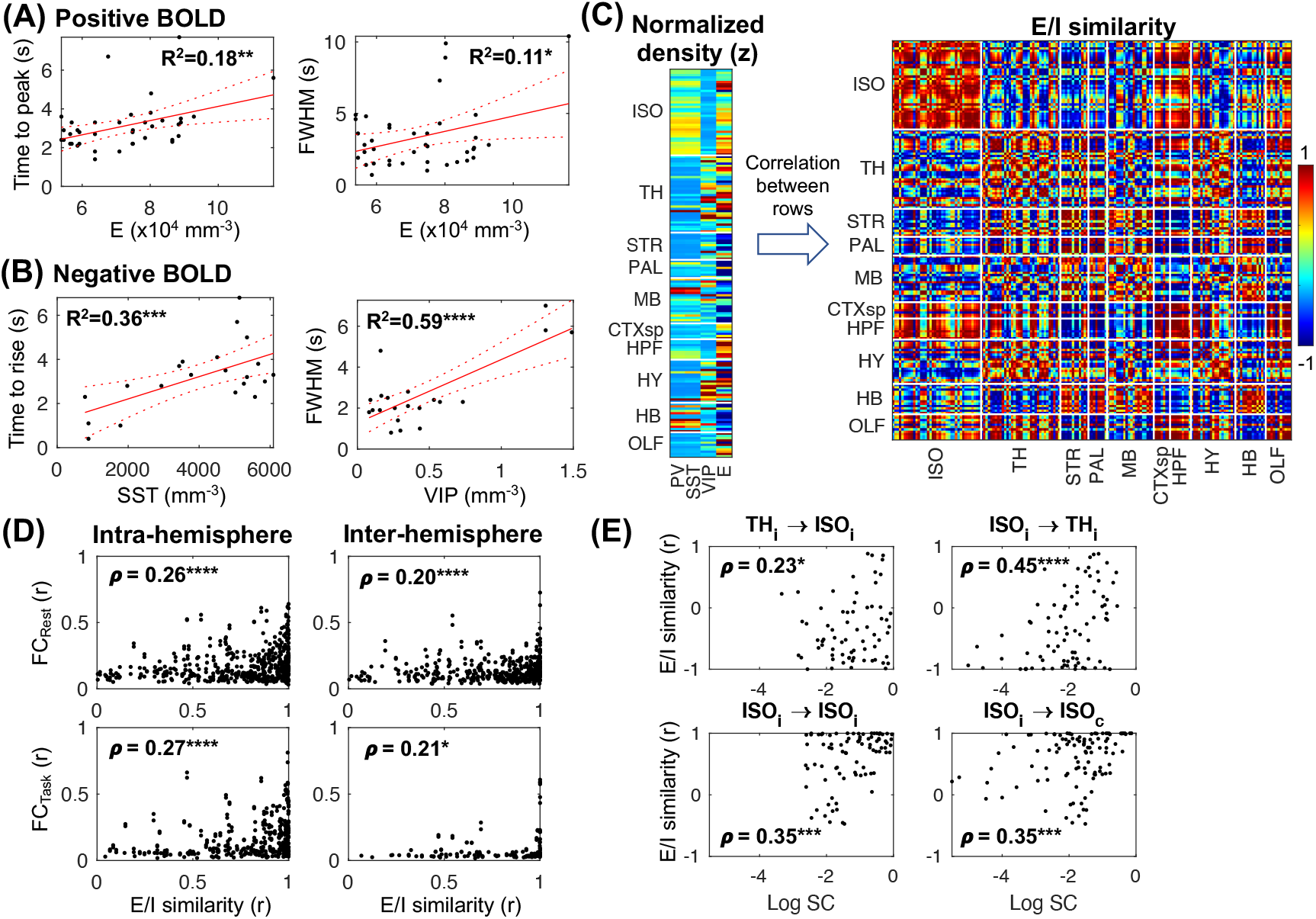
Hemodynamic response, FC and SC are associated with E/I distribution. **(A)** The cortical hemodynamic time-to-peak and FWHM of the positive BOLD correlated with excitatory neuron density. See supplementary Fig. S7 for complete results. **(B)** The cortical hemodynamic time-to-rise and FWHM of the negative BOLD correlated with inhibitory neuron density. **(C)** The normalized densities of 4 types of E/I neurons formed the E/I profile of each region. Inter-regional correlation of the E/I profiles shows the highly similar distribution of the 4 neuron types within the ISO. **(D)** The intra- (left) and inter-hemispheric (right) cortical FC_rest_ (top) and FC_task_ (bottom) correlated with the E/I similarity. **(E)** E/I similarity correlated with SC in both feedforward and feedback projections.

To determine whether the E/I neuronal distribution contributed to the connectivity, we estimated the inter-regional similarity of the E/I distribution by correlating the normalized neuronal density profiles between regions to create an E/I similarity matrix (Fig. 7C). The HPF had the most similar E/I distribution (r=0.89±0.12), followed by the ISO (r=0.63±0.44). In contrast, the TH (r=0.25±0.66), MB (r=0.29±0.63), and STR (r=0.21±0.76) had heterogeneous neuron types. Within the ISO, the E/I similarity within functional modules was generally higher than between modules (supplementary Fig. S9A). For instance, E/I similarity was 0.84±0.16 and 0.93±0.078 within the somatosensory and visual cortices, respectively, whereas the similarity between these regions was 0.65±0.30. The heterogeneity was mainly driven by excitatory neurons as the inter-regional similarity of inhibitory neuronal density was generally high (supplementary Fig. S9B,C).

Compared with the cortical FC (Fig. 7D), there was weak correlation between E/I similarity and FC_Rest_ (ρ=0.26, p=3.8×10^-10^) and FC_Task_ (ρ=0.27, p=2×10^-7^) within hemispheres, and even less between hemispheres (FC_Rest_ ρ=0.20, p=1.5×10^-5^; FC_Task_ ρ=0.21, p=0.013). Interestingly, regions with similar E/I neuronal distribution also tended to have stronger SC (Fig. 7E), including the thalamocortical (ρ=0.23, p=0.039) and corticocortical (ρ=0.35, p=0.00097 intra-hemisphere; ρ=0.35, p=0.00026 inter-hemisphere) feedforward projections, and the corticothalamic feedback (ρ=0.45, p=0.000026). To determine whether the SC or E/I neuronal distribution contributed more to the FC, we used either or both factors as predictors of FC in a linear model (supplementary Table S2). The goodness-of-fit R^2^ to the cortical FC_Rest_ increased when using E/I similarity either alone or together with the SC, whereas the prediction of FC_Task_ only showed marginal improvement. These results indicate that cortical regions of similar neuronal compositions tend to connect with each other. In particular, FC_Rest_ was more dependent on E/I distribution, whereas FC_Task_ more dependent on SC.

## Discussion

This study evaluated two structural constituents, axonal projection and E/I neurons, of the system-level functional dynamics over the hierarchy of a somatosensory pathway. We found that evoked BOLD responses and FC_Task_ correlated with feedforward projection and E/I neuronal distribution. This association decreased with hierarchy due to weaker and more variable neural activation in higher order areas. These characteristics became obscure when the evoked responses were less linear. We also found that hemodynamic timing and shape reflected E/I neuronal density and hierarchy, which could provide information of regional processing and information flow, respectively. In particular, we discovered that cortical regions with similar E/I distributions had stronger SC and FC, with FC_Rest_ corresponding more to E/I similarity and FC_Task_ being more aligned with SC. This greatly expands the findings of previous studies which only examined the first-order projection on the cortical surface (46) or undirected connectivity (10–13). The findings that BOLD transients follow the SC and E/I distribution support the use of hemodynamic signals to probe system-level brain network dynamics and to infer causality. The hierarchy-dependent hemodynamic response indicates a need to refine the current hemodynamic model for data analysis, neural modeling, and interpretation. Our results provide insights into the neural constructs of the structure-function relationship and highlight their differential contributions over the hierarchy and task states.

We demonstrated that ultrafast fMRI allows the detection of brain-wide hemodynamic transients under short stimulus events which improves the characterization of structure-function relationships. Previous fMRI studies mostly detected activation in the ipsilateral somatosensory cortex (47), likely due to the anesthesia effects (48, 49), a weak thalamic response (50) or insufficient sensitivity (31, 51). Limited downstream responses were reported even with direct optogenetic stimulations of the cortex (52) or the thalamus (53). With improved sensitivity provided by a cryogenic coil or ultrahigh field MRI, more activation was recently observed in awake (33) and anesthetized mice (31, 34, 54). Although traditional block design could increase sensitivity, it made responses more nonlinear and harder to associate with the SC.

The weaker and more variable BOLD responses in high-order areas is consistent with an increased activity timescale (55) and variation of latency (56) over the hierarchy instead of baseline CBF or neurovascular coupling. A slower BOLD response in high-order areas accords with the time-to-rise of the LFP as well as recent studies which showed that initial hemodynamic transients can reflect neural information flow in the visual and somatosensory pathways (34, 57, 58). However, the relationship between neuronal and hemodynamic timing remains unclear. We also observed broader FWHM of the LFP at higher levels of the hierarchy. Such hierarchy-dependent dynamics could originate from multi-synaptic connections which lead to a longer “temporal receptive window” in high-order association areas but shorter activity in the sensory cortex (59). This was recently identified in humans using resting-state fMRI (60).

We found distinct hemodynamic responses related to structural organization and hierarchy, with the responses in the STR/PAL being slower than other regions. Such brain structure-dependent hemodynamics could be due to activation of different cell types owing to different projected targets and cellular distributions (61). Specifically, a biphasic response was only seen in the contralateral cortex. Transcallosal interhemispheric inhibition plays an important role in sensory processing, motor control and neuroplasticity (62, 63) by excitatory efferents from the ipsilateral cortex to the local interneurons in the contralateral hemisphere. How inhibitory activity manifests in the BOLD signal is still uncertain. Although a negative BOLD signal would be expected, previous studies also reported a positive response (63, 64). Recent studies found biphasic BOLD or blood flow responses by optogenetic activation of inhibitory neurons, particularly SST^+^ neurons (21, 65). We also found that a negative BOLD response associated with inhibitory neuronal densities. Together, these findings indicate that a negative BOLD signal reflects transcallosal inhibition. A previous study showed bilateral positive activation under VPM excitation in isoflurane-anesthetized rats (40). Compared to the medetomidine used in this study, the hemodynamic response under isoflurane is generally unipolar (66). This is likely due to isoflurane-induced disruption of E/I balance (67) and functional hyperemia mediated by inhibitory interneurons (22).

Both evoked activation and FC correlated with feedforward, but less with feedback, projections. Although this may be due to the suppression of cortical feedback by anesthesia (68), a study in awake monkeys found that thalamocortical feedforward spiking activity increases with stimulus intensity whereas corticothalamic feedback remains unchanged (69). This may be due to the interaction of multiple sources of feedback information and the function of a feedback route. Although corticothalamic projections are mostly excitatory, they could also induce inhibition via the RT (70). In the sensory pathway, feedback signals have diverse modulatory roles in behaviors such as attention (71). Further differentiation of excitatory versus inhibitory connections will help to clarify the weak coupling in feedback projections.

We found that E/I ratio increased over the hierarchy and may underlie the increased BOLD variability. This is consistent with a macroscopic gradient of E/I distribution and top-down control of sensory inputs (25). We also found that E/I similarity correlated with both structural and functional connectivity and was a major contributor of the FC_Rest_. Incorporating this information may improve the modeling and prediction of neural dynamics. Why regions with similar E/I distribution have stronger SC and FC is unclear. This may be due to similar genetic profiles and developmental origins. For instance, cortical excitatory neurons that originate from the same progenitor cell have stronger synaptic connections, share similar stimulus selectivity and form long-range connections with the same microcircuit in high-order areas (72, 73).

There are several limitations of this study. Although we used one of the most reliable anesthesia protocols for mouse fMRI, the anesthesia could affect both neural and hemodynamic responses in pathways involving thalamic and limbic areas (74). This issue could be minimized by further development of awake fMRI that incorporates optogenetics (75). Secondly, we activated all neurons to allow comparison with the SC that does not differentiate cell types. As different types of neurons have preferential projections, activated regions can be cell-type dependent (61). Further study using cell-type-specific projections and stimulations would allow the delineation of a more precise relationship in excitatory and inhibitory connections. Thirdly, besides axonal and E/I neuronal densities, other factors such as neuromodulators, synaptic density and receptor subtypes, have been found to change over the hierarchy (23, 24). To verify their specific contributions, further studies involving inhibition of a specific cell type, receptor or projection using optogenetics (76) or chemogenetics (77) will be needed. Fourthly, the interhemispheric FC_Task_ was underestimated due to the distinct hemodynamic response in the contralateral cortex. HRF deconvolution may improve the estimation but this requires a high signal-to-noise ratio. Finally, due to the limited spatial resolution, cortical layers were not differentiable. Pushing the spatial resolution using higher field MRI will enable comparison with laminar input-output projections (76) and E/I neuronal distribution to further understand the structure-function relationship at the mesoscale.

## Materials and Methods

Animal experiments were approved by the Animal Ethics Committee of the University of Queensland. Four groups of male C57BL6/J mice were used: one to evaluate the heating effects of optogenetic stimulation (n=6 with ChR2 and n=3 without virus), the second (n=8) to investigate responses to different stimulus intensities and frequencies, the third (n=6) to measure electrophysiology, and the fourth as a naïve group without surgery to acquire resting-state fMRI (n=11) and CBF (n=4). The mice were aged 6-7 weeks at the time of viral injection and 9-15 weeks at the time of fMRI. The details of animal preparation, experiments, and data analysis are provided in the SI Appendix, Supplementary Methods.

## Acknowledgments

We thank Prof Linda Richards for helpful discussion and Ms Rowan Tweedale for proof reading. H. H.H. was funded by the Australian Research Council grant number DP180103319. K.H.C was supported by startup funds from the Queensland Brain Institute and Centre for Advanced Imaging, The University of Queensland.

